# Spatial structure imposes sex-specific costs but does not reduce interlocus sexual conflict

**DOI:** 10.1101/2022.10.29.514349

**Authors:** Subhasish Halder, Shramana Kar, Simran Sethi, Swadha Tewari, Tanya Verma, Bodhisatta Nandy

## Abstract

Spatial structure is a common feature of all naturally occurring populations. Theoretically, spatial structuring of a habitat could modulate the intensity of Interlocus sexual conflict (ISC) in a population, possibly by modulating intersexual encounter rate. We tested this theory using laboratory populations of *Drosophila melanogaster* by measuring male induced fitness decline in females in three-patch habitat systems under two alternative habitat types – structured-interconnected and unstructured. Our results on reproductive and survival costs in females suggested significant costs due to (a) male presence (i.e., ISC) and (b) living on structured habitat. However, there was only a weak evidence supporting the theory of modulation of ISC by habitat structuring only. Through a follow up experiment, we further showed that the effect of habitat on ISC is also robust to the alteration of female conditions. Therefore, it appears that spatial structuring per se is unlikely to modulate ISC, but can impose additional survival costs. We further showed such survival cost could be sex-specific possibly reflecting female biased spontaneous dispersal.

## Introduction

In absence of a true monogamy, males can maximize their fitness by investing in competitive and manipulative traits that increase their mating and/or fertilization success. In contrast, female fitness is usually maximised by achieving longer lifespan, higher fecundity with better-quality eggs, and ensuring higher survival rate of juveniles (Holland & Rice, 1999; Arnqvist & Rowe, 2013; Adler & Bonduriansky, 2014). Such asymmetry in evolutionary interests across the two sexes is commonly referred to as sexual conflict, which is now recognized as a widespread phenomenon affecting many animal and plant taxa (Parker, 1979). Theories and empirical evidence suggest that male-benefiting traits that result in better mating and/or fertilisation success can emerge as male reproductive adaptations regardless of their impact on females (Rice, 1996; Chapman et al., 2003; Nandy et al., 2013). Often, such male-benefiting traits are detrimental to female fitness as they involve coercing females to mating and/or reproducing at a time and rate which benefits males but is injurious for long-term survival of the females, thus reducing the lifetime progeny output of the females. This sets the stage for the emergence of female counter-adaptations that minimise female susceptibility to male manipulations (Chapman et al., 2003; Wigby & Chapman, 2004; Nandy et al., 2014).This form of sexual conflict, commonly referred to as interlocus conflict, can result in intersexual antagonistic co- evolutionary armsrace (Chapman et al. 2003; Pizzari & Snook 2007).

A crucial aspect of interlocus conflict is the involvement of direct interaction between the sexes such as, persistent and coercive courtship, forced copulation, traumatic insemination, males riding the females or guarding them for long duration, and even modulation of female physiology by the seminal fluid proteins and peptides transferred during mating (Arnqvist, 1992; Chapman et al., 1995; Réale et al., 1996; Le Galliard et al., 2005; Adller, 2010; Koene, 2012). In fruit flies, *Drosophila melanogaster*, females suffer increased mortality and/or reduced life-time progeny production due to coercive male mating behaviour and toxic seminal fluid peptides received during copulation (Chapman et al., 1995; Rice et al., 2006; Wolfner, 2009; Nandy et al., 2013). Most theoretical and empirical investigations of interlocus conflict that deal with the intensity of conflict in a population tend to assume, either implicitly or explicitly, an unbridled interaction between the sexes that allow the conflict to operate (Rice et al., 2006). However, most natural populations are spatially structured, and hence, individuals are able to move around across patches seeking or avoiding favourable and unfavourable interactions. The way such population structure affect intensity of conflict in a population is poorly understood.

It is not unreasonable to expect females to avoid patches with a high male congregation to avoid mate harassment. An explicit experimental verification of this theory was first attempted by Rice et al., (2006), wherein *D. melanogaster* females were found to suffer less male harassment (measured as remating rate) when they were allowed to move between two compartments within a culture vial.

However, subsequently, more thorough work from the same group did not find the intensity of sexual conflict to be lower when the experimental habitat allowed spatial refuges (Byrne et al., 2008). More recent experiments on *D. melanogaster* showed that the intensity of interlocus conflict is significantly reduced in a large and heterogenous holding environment compared to a small and homogenous one (Yun et al., 2017; Malek and Long, 2018). In addition, the scope of hitherto reported adaptive male mating bias, a source of mate harassment, in favour of high fecundity females appeared to have reduced in the large and complex environment (Yun et al., 2018). In water striders of the species *Aquarius remigis*, physical structuring of population modulates males’ aggressive behaviour towards females (Eldakar, 2009).

Further, individuals in a patch structured population are expected to disperse across population patches, resulting in an added complication due to the ecology of dispersal. Indeed, in *Aquarius remigis*, maintained in a naturalistic laboratory environment that introduces dispersal alters male-female local interactions thereby altering the dynamics of interlocus sexual conflict. For instance, in isolated pools, without dispersal, males tend to gain more mating by being aggressive. Interestingly, in interconnected pools which allowed individuals to disperse, females were found to disperse away to avoid aggressive male, which in turn affected male dispersal (Eldakar, 2009). Besides, if there is inherent difference in dispersal tendency across two sexes, which is prominent in many animals (Trochet, 2016; Mishra et al., 2018a), it can heavily influence male-female interactions, thus shaping interlocus conflict and its evolutionary consequences. In addition, it is important to note that dispersal may also impose novel ecological costs. For example, dispersal and other movement related traits such as foraging or exploration, are energetically and/or ecologically costly (Bonte et al., 2012). However, it is not clear how such costs and benefits affect the outcome of interlocus conflict in a population. MacPherson et al., (2018) found low condition females, raised in high larval density, suffered from the cost of increased male exposure, regardless of their holding condition, whereas high condition females were found to be practically immune to mate harm in complex, but not in simple habitat. A patch structured habitat, apart from being relatively complex, may bring in an additional ecological component of dispersal. If both dispersal and resistance to mate harm are energetically expensive, structuring of habitat can impose additional cost on females thereby making them more susceptible to mate harm. However, this expectation will depend on cost of dispersal as well as female condition. Therefore, spatial structuring of a population can clearly impacts (a) the level of interlocus conflict in a population, and (b) the evolutionary trajectory of sexually antagonistic traits. However, more investigations are needed to test the theory and assess the general implications. With ever increasing anthropogenic fragmentation of natural populations with varying degrees of connectivity within patches (Rogan & Lacher Jr., 2018), it is becoming increasingly more important to investigate the impact of such changes in the habitat structure on key eco-evolutionary processes, such as, sexual conflict.

Here, we addressed this issue using a *D. melanogaster* laboratory adapted populations. We investigated whether spatial structure could also have a similar complex habitat effect of relaxing intensity of sexual conflict. We mimicked spatial structure by setting up three-patch habitat systems by connecting three standard culture vials with narrow tubes that allow flies to move from one patch (vial) to another. As controls, three unconnected vials represented similar interacting population units, but not distributed following any spatial structure. The control habitat is similar to the usual laboratory population, without a spatial structure wherein the sexes are free to interact with each other with no scope of dispersing to a different habitat patch, and without an access to spatial refuges. The experimental structured habitat type retained the possibility of identical interaction arena but also allowed individuals to move across patches (i.e., vials) and potentially avoid antagonistic male interactions. Further, individuals within a patch in such a setup, have a higher chance of interacting with each other compared to those from two different patches. We then compared two components of female fitness - fecundity and starvation survival time, following two types of male exposures – limited exposure and continuous exposure, across these two habitat treatment types. We used the results to test the hypotheses on cost of male exposure in females (i.e., the reproductive cost in females arising due to ISC) and cost of dispersal. The benign and nutritionally rich laboratory maintenance regime of populations such as, the ones used in our experiments, can often obscure fitness costs, especially those concerning resource allocation trade-offs (Van Noordwijk & De Jong, 1986; McCracken et al., 2020; MacTavish & Anderson, 2020).

Therefore, we further investigated this issue using resource deprived females. To this effect, we first investigated the role of developmental dietary restriction on spontaneous dispersal tendency to determine a level of dietary manipulation that does not significantly alter dispersal but results in significantly resource deprived adults. We then repeated the above-mentioned experiment with an added treatment of the female diet regime wherein standard and nutritionally deprived females were used.

## Material and methods

We used flies from a set of large, outbred, wild-type, laboratory adapted populations of *Drosophila melanogaster* (BL_1-5_). In 2010, 18 iso-female lines generated from wild caught females in Blue Ridge, USA, were mixed to create a mass-bred population. Five replicate populations were established by collecting a random set of eggs ∼2,500 eggs. These replicate populations (BRB populations) were brought to the lab in 2015 to establish the five BL populations. Hence, these populations can be viewed as replicate outbred populations generated by random sampling from a single global population.

These populations are maintained under a 14-daydiscrete generation cycle, 24 h light, at 25 °C (±1) temperature on standard banana-jaggery-yeast food medium. Flies are grown in culture vials (25 mm diameter × 96 mm height) at a density of ∼70 per 6-8 ml food in each vial, forty such vials make up one population. On day 12 post-egg collection, and after all the adults have emerged from pupae, flies are transferred into a population cage and thereafter maintained in the cage as a population of ∼2,800 individuals. On day 14, eggs are collected on a fresh food plate within a time window of 18 h. These eggs are cultured in fresh food vials following the above-mentioned density to start the next generation. Therefore, the progeny produced during this 18 h window can be considered total lifetime progeny output, and an accurate measure of Darwinian fitness (i.e., direct component of fitness) in this system (see Rice et al. 2006). The details of the history and maintenance of the populations can be found in Nandy et al., 2016 and Dasgupta et al., 2022. Three randomly chosen BL populations, viz., BL_2_, Bl_3_ and BL_4_ were used to conduct the experiments described below.

Structured and unstructured habitats and the experimental setup:

As mentioned above, the base populations are maintained as large unstructured island populations, either as a single large unit with ∼2,800 individuals (i.e., the cage phase), or 40 small unconnected units each with ∼70 individuals (i.e., the vial phase). For the purpose of the experiments, we used a simple three-patch setup. We created experimental patch-structured habitat systems (hereafter simply referred to as “structured” habitat type), by connecting three culture vials with narrow tubes (length 12.5cm, diameter 0.6 cm, hereafter referred as “corridor”) that allowed flies to move from one vial to another in either direction (see Fig. S1). Results from pilot studies conducted prior to the assays suggested that both density of flies (individuals per patch) and sex ratio in each patch show substantial spatial and temporal variation (Figure S2a and S2b) within the framework of the assay conditions mentioned below. Further details regarding the dispersal behaviour of flies held in such an experimental setup can be found in the Supplementary information.

We used three standard culture vials, without connection between them, as controls (Fig. S1). Hereafter, we refer to this habitat type as “unstructured”). We opted for this design, since we aimed to compare structured vs. unstructured habitats controlling for the population size (i.e., large vs. small) effect. In the structured setup, because individuals were free to move around, the density across the three patches are expected to change (also see SI, Figure S2a). However, the average density (number of individuals per vial) was expected to remain the same between the two habitat types.

### Experiment 1

In the first experiment (Experiment 1), we investigated the effect of habitat structure on the extent of interlocus conflict. We measured progeny production in females under two types of male exposures – limited exposure (LE: females exposed to males for a short duration that ensured only single mating) and continuous exposure (CE: females continuously held with males). The effect of male exposure type treatment on female progeny output (a good measure of female fitness) in such an assay has been found to be a good measure of interlocus conflict in similar *D. melanogaster* systems (Nandy et al. 2013, 2014). The assay was done under two habitat types – structured and unstructured. In addition to progeny production, we also measured starvation survival time as a measure of the survival component of fitness that is often affected by dispersal and reproduction (Salmon et al., 2001; Flatt, 2011; Harvanek et al., 2017; Mishra et al., 2022). The entire experiment was performed in a randomized block design with three independent statistical blocks using three randomly chosen replicate base populations (viz., BL_2_, Bl_3_ and BL_4_).

To generate experimental flies, eggs were collected from the baseline populations in standard food vials (∼8ml food) with a density of ∼70 eggs/vial. After egg collection, vials were kept in controlled laboratory condition and monitored carefully for the onset of emergence of adults. Following the onset of eclosion, flies were collected every six hours. Experimental flies were collected as virgins from the peak of eclosion during which flies were separated by sex and housed in same sex vials at a density of 5 females/vial and 10 males/vial. The assays were performed on 2-3 days (post-eclosion) old flies. This adult age was chosen because it is the most relevant to adult fitness in this system – viz., 2-4 days post-eclosion age corresponds to height of adult interaction and progeny production window.

On day 12, 10 virgin males and 5 virgin females were introduced in each vial for both S and US setups. An adult sex ratio of 2:1 was taken to ensure higher male inflicted harm on females, thereby helping in the resolution of even relatively smaller differences in female performance across treatments. For the unstructured habitat type, a set of three identical vials with 10 males and 5 females in each vial (i.e., 30 males and 15 females for a set) constituted a replicate set. For the structured habitat type, the same numbers of sexes (i.e., 10 males and 5 females) were introduced in each of the three interconnected vials. The cotton plugs of each vials were pushed to half to account for the standard population density as number of flies/space. Twenty-four replicate sets for each of the unstructured and structured habitat types were established. These twenty-four replicates were further subdivided into limited exposure (LE) and continuous exposure (CE), therefore twelve replicate sets in each male exposure treatment. Thus, our assay design involved a 2 X 2 fully orthogonal combination of the two treatments (viz., habitat and male exposure type). Therefore, we had 12 replicates for each male exposure × habitat type treatment. For the LE subset, one hour following the introduction of experimental flies, males were removed (using light CO_2_ anaesthesia) whereas females were retained in the same setup. Our personal observation suggests the majority of virgin females get their first meeting within this one hour. In the CE subset, males and females were allowed to remain in their respective setups for ∼48 hours. To equalise the handling, flies in the CE setup were also exposed to mild dosage of CO_2_ roughly after 1 hour after the initial introduction of the sexes in the setups. On day14, i.e., following ∼48 hours of the initial combination of the sexes, flies were anaesthetized and ten randomly chosen females were transferred to oviposition tubes. For both unstructured and structured setups, females from the three component vials in a replicate setup were combined and 10 females (out of 15) were randomly chosen for oviposition. Males were discarded at this point. A fixed window of 18 hours was allowed to the females in the oviposition tubes (1 female in each oviposition tube) for laying eggs. Following this, the females were removed from the tubes; the tubes were then incubated for offspring emergence. Per capita progeny output was calculated for a given replicate as the average number of offspring produced by these ten females. Following oviposition in tubes, 5 females were randomly chosen out of 10 and were regrouped and transferred into a non-nutritive agar vial to measure starvation survival time.

Female survival in these vials were recorded every six hours. Mean survival time was calculated as the average survival time of five females for a given replicate vial.

### Experiment 2

Dispersal tendency and susceptibility to mate harm in females are known to be condition dependent (Van Noordwijk & De Jong, 1986; McCracken et al., 2020; MacTavish & Anderson, 2020). Relatively rich nutritional environment of our standard laboratory regime can obscure the fitness costs that depend on resource allocation trade-offs. Therefore, we repeated the above experiment with an added treatment - dietary condition. However, *a priori* information was needed on female response to nutritional manipulations. Hence, we first conducted an assay to measure the effect of manipulation of developmental diet on female dispersal tendency (see supplementary material). The results suggested that a nutritionally diluted developmental diet, even up to 40% reduction compared to the standard diet, does not affect female dispersal tendency (Fig. S6). Part of these results however, also suggested that such dietary treatment significantly reduces female quality as measured by lower body mass at eclosion (also see Poças et al., 2022). Such females are considered resource deprived or poor condition females.

We conducted Experiment 2 with poor and standard condition females. Additionally, in this experiment, we also assessed the cost of dispersal for males. The design of this experiment was similar to that followed in Experiment 1, except the additional treatment for female condition. We use nutritionally challenged females (poor condition: larval development in 40% diluted food) and those raised on standard food (control: larval development in standard banana-jaggery-barley-yeast food). The detailed composition of the food types is mentioned in the supplementary information (Table S2).

Flies were generated using a method similar to that followed in Experiment 1. However, as mentioned above, for this experiment, two types of females - poor and standard, were raised and collected as virgins (see Experiment 1). Followed by this, the experimental flies were subjected to the assay setups. However, unlike Experiment 1, this was done on 3-4 days old flies to account for the difference in development time of the poor and control females. We subjected the females to two habitat treatments - unstructured and structured, as mentioned in Experiment 1. These setups were further subdivided into two mate exposure treatments - limited exposure and continuous exposure, i.e., the LE and CE vials as mentioned in Experiment 1. All males used in this stage were raised on a standard diet. Subsequently, we measured female progeny output and starvation survival time following the protocol similar to that followed in Experiment 1. We had 8 replicate setups for each treatment and number flies in each vial/setup were identical across Experiment 1 and 2. Experiment 2 was also performed in randomised block design with three blocks by using three randomly chosen BL populations - BL_3_, Bl_4_ and BL_5_. Per capita progeny output and mean starvation survival time for the experimental females were computed following the same method adopted in Experiment 1.

### Statistical analysis

All experiments were conducted following a randomized block design. Assays were repeated in three randomly chosen replicate populations. It is important to note that these populations represent a sample of outbred laboratory adapted populations from an infinite number of such populations that could be created by randomly sampling from the natural population of *D. melanogaster.* Hence, population (or block) was treated as random factor. Our interest was to test the effects of the other fixed factors controlling for the random effect of the block. All treatments within a given population were handled together and the assays on three replicate populations were done on three different days. Hence, even by handling, populations were treated as random blocks. Therefore, despite the recent debate over mixed model analysis when random factor has less than five levels (Bolker, 2015), we treated population identity (i.e., block) as a random factor. Progeny count and mean starvation survival time results were analysed using a linear mixed-effect model using `lmer` function of the `lme4` package in R (R Version 4.2.0). In the models, habitat type, male exposure type, and female type (only for Experiment 2) were treated as fixed factors, while block and all interactions concerning block were modelled as random factors. In order to test for the effect of random factors we ran separate models by dropping a random factor term of our interest, followed by model comparisons with the global model using `anova` function. Pairwise multiple comparisons were done with `pairwise` function in package `emmeans`. We also estimated effect size (Cohen’s *d*) for the statistically significant effects using package `effsize` and function `cohen.d`. Following Cohen (2013), the effect size is interpreted as small (*d* < 0.5), medium (0.5 < *d* < 0.8) or large (*d* > 0.8). In addition to the mixed model analyses as stated above, we re-analysed the experimental results using purely fixed effect models. The methods and results of these analyses are mentioned in the supplementary information. All our conclusions are robust to the treatment of block as random or fixed factor.

## Results

### Experiment 1

Analysis of the progeny count indicated that effects of both male exposure and habitat type treatment were significant (Table 1). On an average, progeny count from females of the LE treatment was 55% higher compared to that of the CE treatment (Fig. 1a, Cohen’s *d* estimate 0.99 with 95% CI: lower CI: 1.35, upper CI: 0.63), indicating a significant fitness cost due to continued exposure to the males.

**Figure 1:**
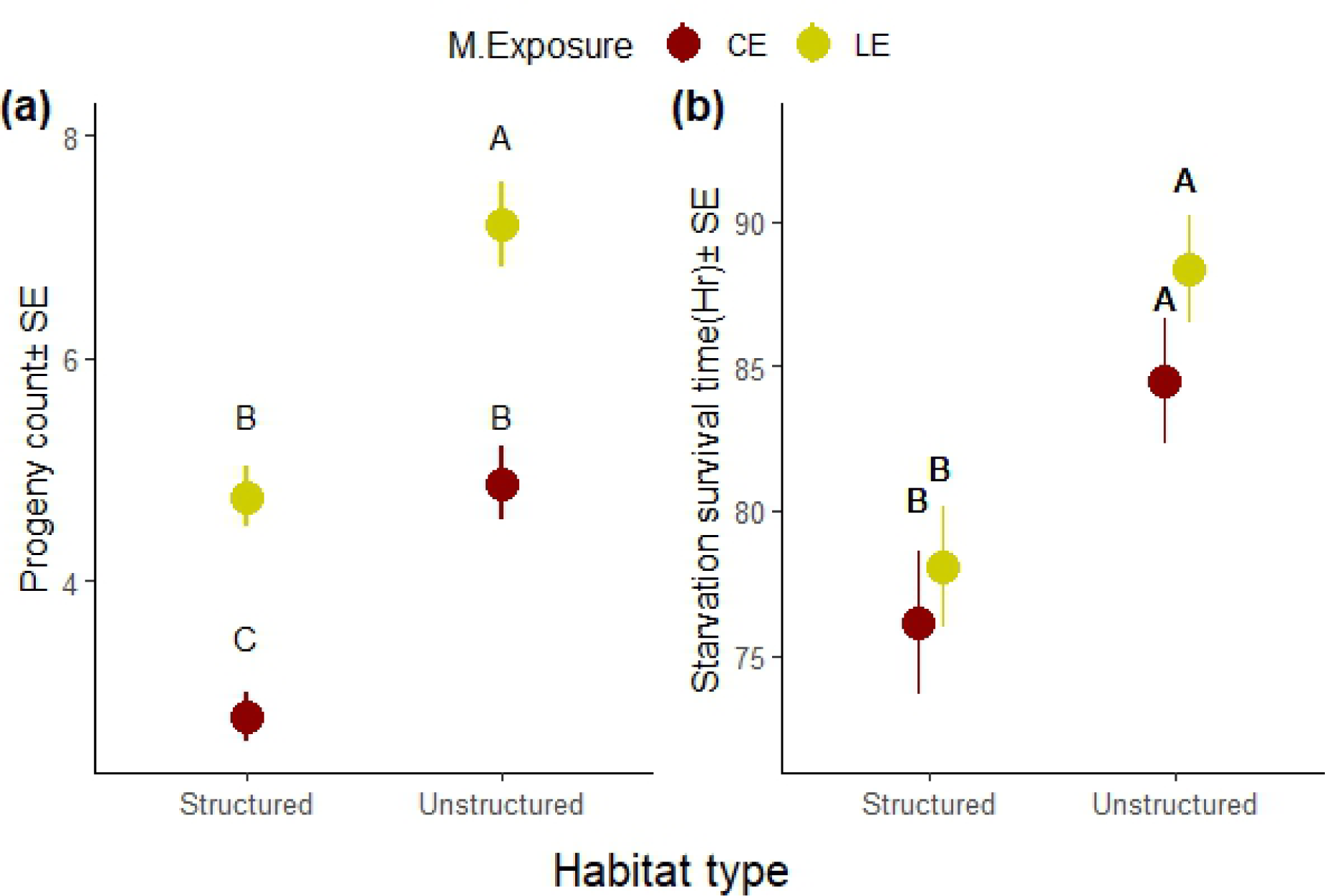
Results from Experiment 1. (a) Progeny count, i.e., number of offspring produced by the experimental females during the 18 h window following the experimental treatment. Per capita progeny output was computed as the average progeny output from ten of the fifteen females in a replicate setup. Per capita progeny count was used as the unit of analysis. (b) Starvation survival time of the experimental females following the treatment and progeny production. Starvation survival time was computed as the mean of the five randomly selected females within a replicate set. This mean was used as the unit of analysis. The treatment represents a full factorial combination of male exposure (limited exposure, LE and continuously exposed, CE) and habitat (structured and unstructured). The plot represents mean progeny output ±SE (a) and mean starvation survival time ±SE (b) of all three statistical blocks. Treatments not sharing common letters are significantly different.

**Table 1:**
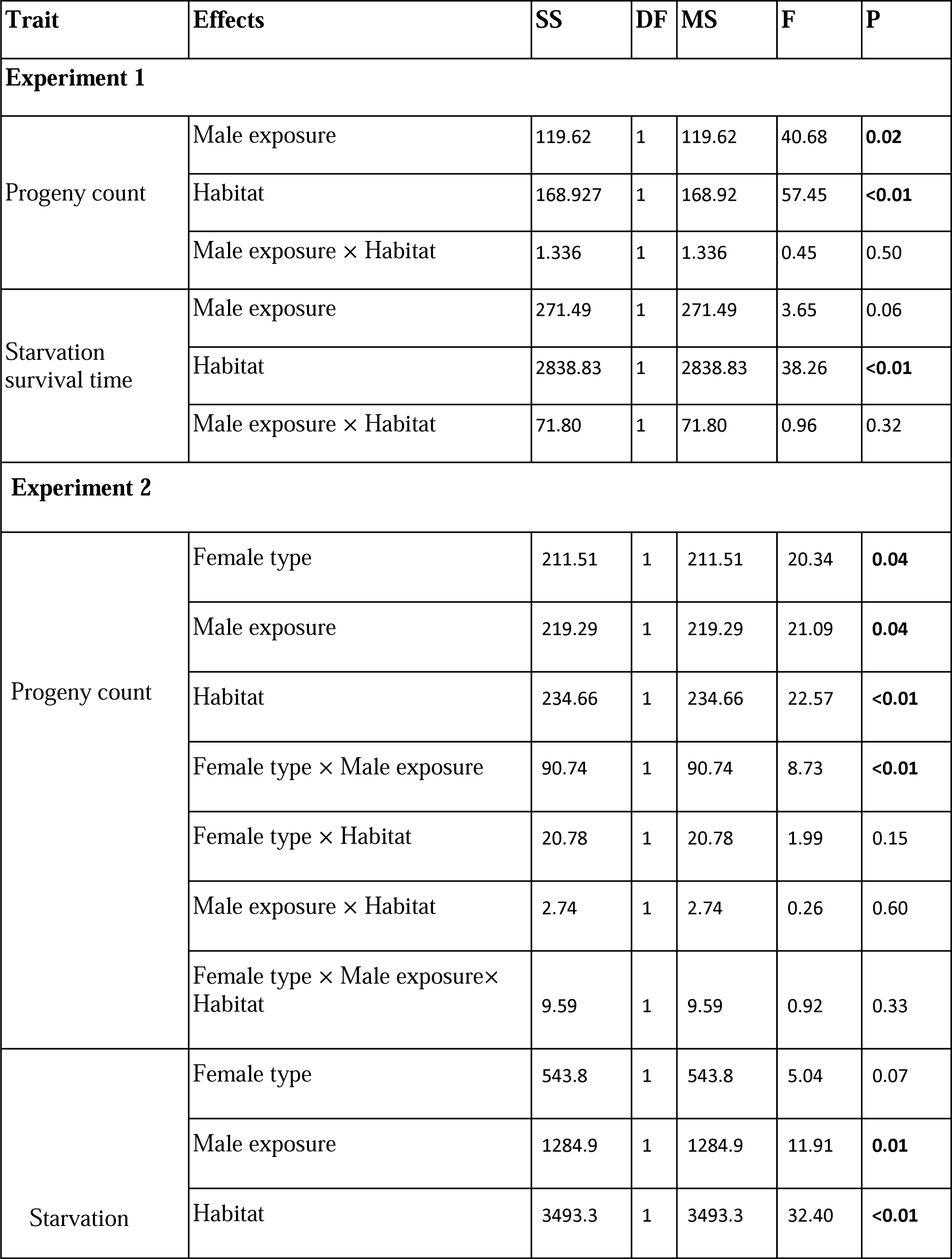
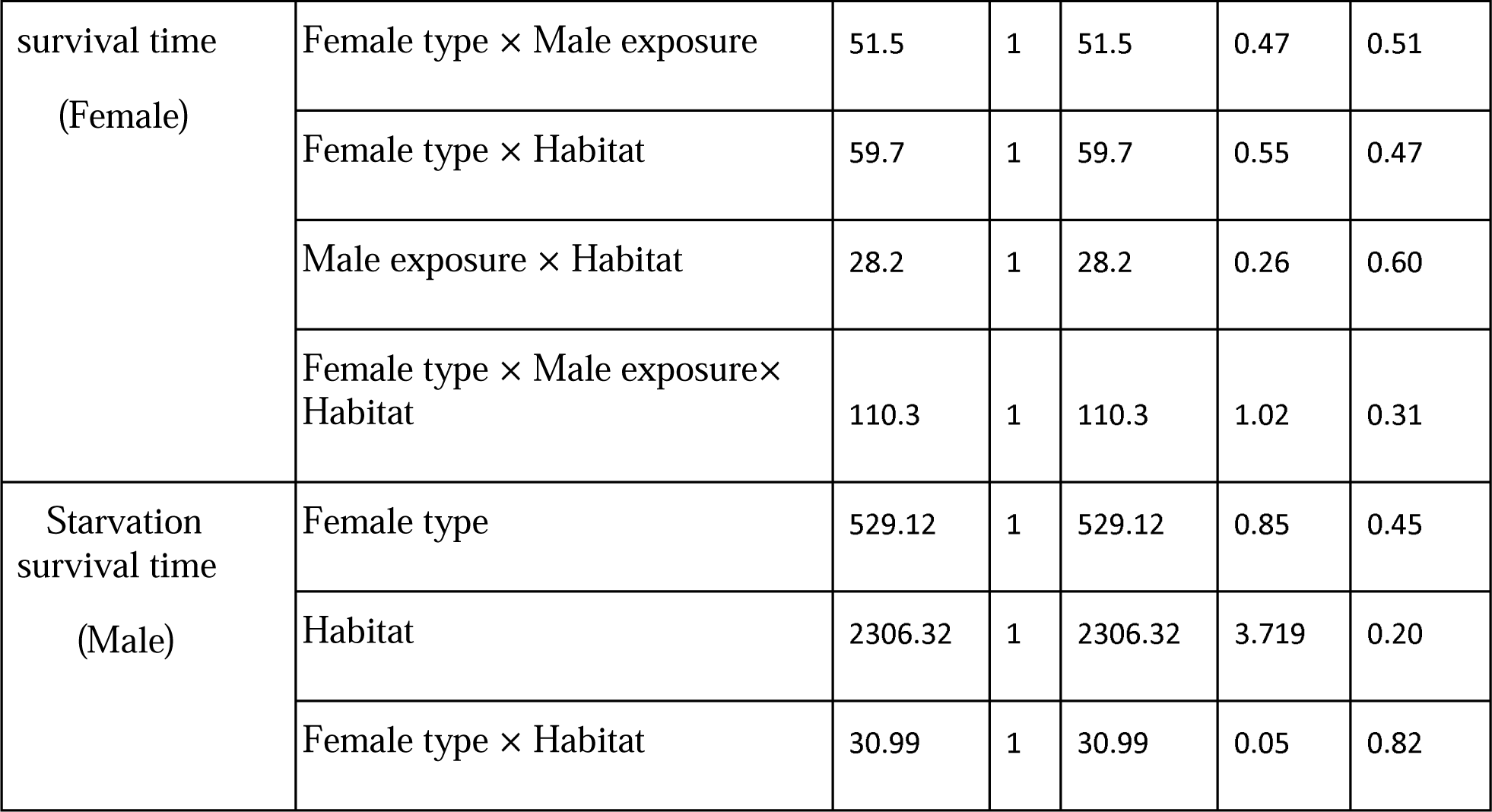
Summary of the analyses of results from Experiment 1 and 2. The results were analysed using linear mixed-effect models where male exposure, habitat type, female type (for Experiment 2) was fitted as fixed factors. Block, and all interactions that included it were fitted as random factors. Statistically significant p-values are mentioned in bold font style.

Similarly, the progeny output of females from structured setup was on an average 37% lower compared to those from the unstructured setup (Fig. 1a, Cohen’s *d* estimate 1.06 with 95% CI: lower CI: 1.43, upper CI: 0.70). However, male exposure × habitat type interaction had no significant effect on progeny count (Table 1). Hence, there was no evidence to suggest that the degree of ISC was affected by the habitat type treatment. Neither the effect of block nor any interaction term involving block on progeny count was found to be significant.

Analysis of the starvation survival time indicated that the effect of male exposure was marginally non-significant (Table 1). Habitat was found to have a significant effect (Table 1), wherein starvation survival time of females from structured treatment was 11% less compared to that of the females from unstructured treatment (Fig. 1b, Cohen’s *d* estimate 0.74 with 95% CI: lower CI: 1.09, upper CI: 0.39) – an indication of survival cost of spontaneous dispersal exhibited by the females in structured habitat (see supplementary information for observation on dispersal across patches). The effect of male exposure × habitat interaction was not significant (Table 1). Neither the effect of block nor any interaction term involving block on starvation survival time was found to be significant.

### Experiment 2

Results of analysis on progeny count indicated significant effects of all three main effects, viz., female type, male exposure and habitat (Table 1). As expected, poor condition females (i.e., females raised on 40% diluted food) produced, on an average, 42% less progeny than standard females (Fig. 2, Cohen’s *d* estimate 0.84, with 95% CI: lower CI: 1.14, upper CI: 0.54). On an average, progeny count of females from the structured setup was on average 29% less compared to those from the unstructured setup, indicating cost of dispersal that comes from living in a structured habitat (Fig. 2, Cohen’s *d* estimate 0.50, with 95% CI: lower CI: 0.80, upper CI: 0.21). Progeny count from females belonging to the LE treatment was on an average 65% higher compared to that from the CE treatment, replicating the finding from Experiment 1, indicating significant fitness cost of male exposure (Fig. 2, Cohen’s *d* estimate 0.77, with 95% CI: lower CI: 1.07, upper CI: 0.47). Interestingly, female type × male exposure interaction was found to have significant effect on progeny count. This suggested that the reproductive cost of male exposure, the measure of ISC, is dependent on female type. Pairwise comparisons (Table S4) revealed a significant reduction of progeny count in CE treatment compared to the LE treatment only for standard females, but not for females raised in poor diet (i.e., poor female type).

**Figure 2:**
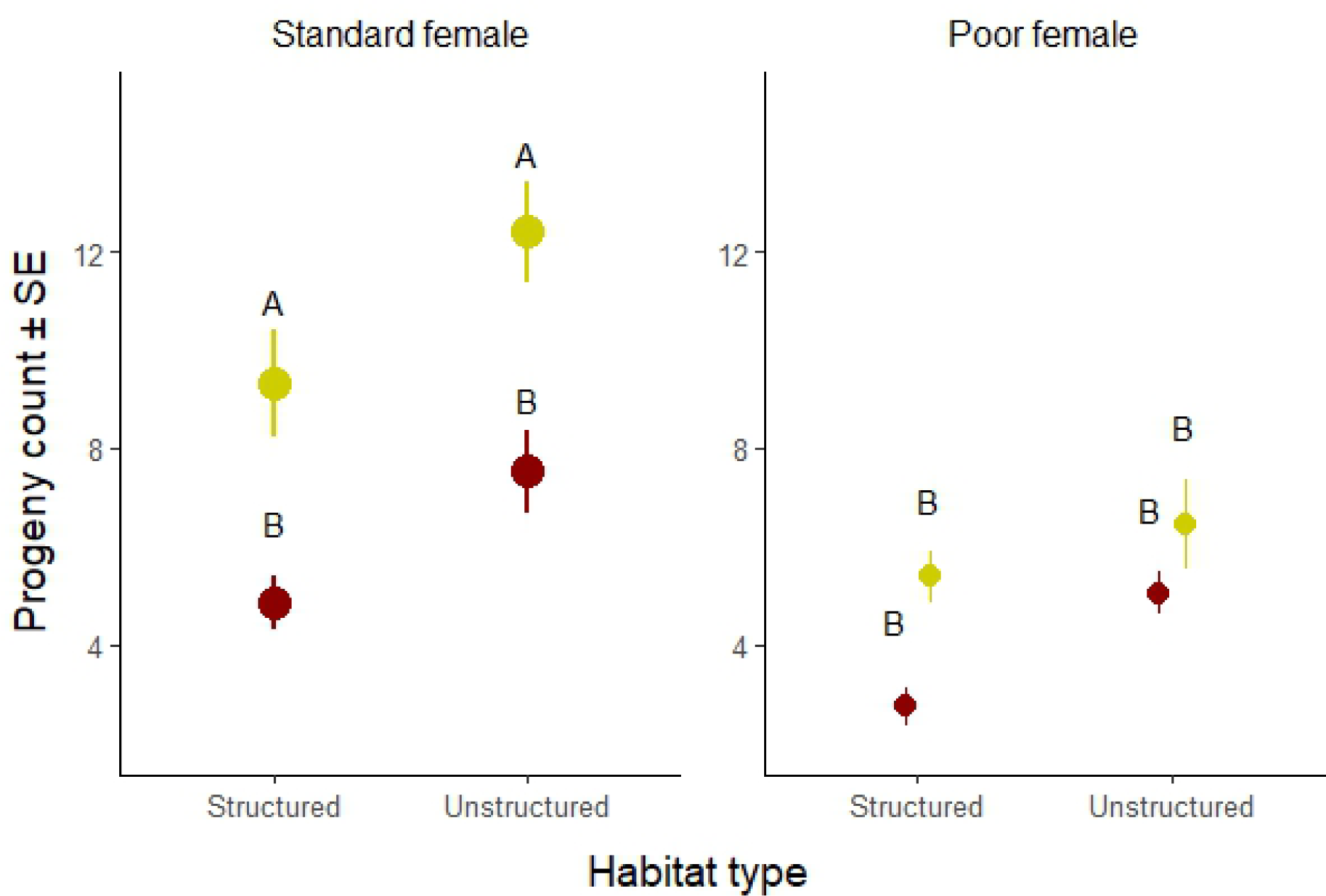
Progeny count results from Experiment 2. Progeny count, i.e., number of offspring produced by the experimental females during the 18 h window following the experimental treatment. Per capita progeny output was computed as the average progeny output from ten of the fifteen females in a replicate setup. Per capita progeny count was used as the unit of analysis. The treatment represents a full factorial combination of female type (poor and standard), mate exposure (limited exposure, LE and continuously exposed, CE) and habitat type (structured and unstructured). The box plots represent data from all three statistical blocks. Treatments not sharing common letters are significantly different.

However, the effects of female type × habitat interaction as well as the male exposure × habitat interaction were not significant (Table 1). Therefore, similar to the results of the Experiment 1, the analysis did not reveal any evidence suggesting that the male induced fitness reduction is affected by habitat type treatment.

Analysis of the starvation survival time of the females revealed significant main effects of male exposure and habitat (Table 1). Effect of female type was marginally non-significant (Table 1, Fig. 3a). Starvation survival time of females from structured treatment was on an average 12% less compared to that of the females from unstructured treatment (Fig. 3b, Cohen’s *d* estimate 0.97, with 95% CI: lower CI: 1.28, upper CI: 0.66), replicating the results from Experiment 1. Females that received continuous male exposure (i.e., CE) died under starvation on an average 6% faster than females that received limited male exposure (Fig. 3c). None of the two-way and three-way interactions was found to have a significant effect on starvation survival time of the females (Table 1). For males, neither the main effects (viz., female type and habitat) nor the interaction between them had a significant effect on starvation survival time (Table 1, mean ±SE: Unstructured: 64.01±7.53, Structured: 78.34±8.29 and Standard: 63.79±6.82, Poor: 78.22±8.80).

**Figure 3:**
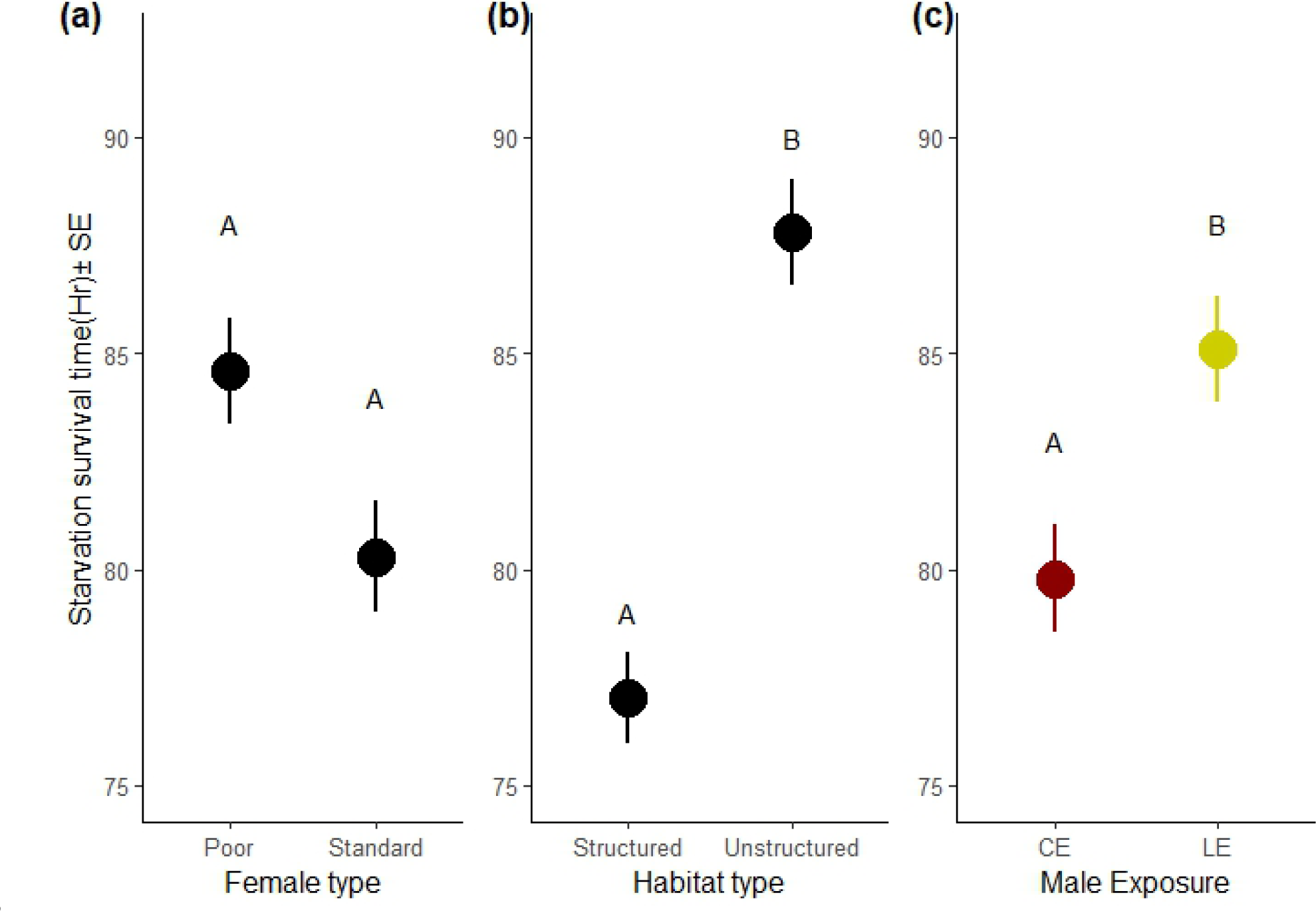
Female starvation survival time results from Experiment 2. Starvation survival time of the experimental females following the treatment and progeny production. Starvation survival time was computed as the mean of the five randomly selected females within a replicate set. This mean was used as the unit of analysis. The treatment represents a full factorial combination of female type (poor and standard), male exposure (limited exposure, LE and continuously exposed, CE) and habitat (structured and unstructured). The box plots represent data from all three statistical blocks. The figure shows the main effects of female type (a), habitat (b), and male exposure (c) on starvation survival time of the experimental females. Treatments not sharing common letters are significantly different.

## Discussion

In both our experiments, regardless of the experimental habitat type we found females to bear a substantial fitness cost of continuous exposure to males. We found such a cost to be expressed in the form of reduced reproductive output, and also as reduced starvation resistance. Importantly, given their laboratory ecology, these costs are good measures of the actual fitness cost – i.e., gender load imposed through interlocus sexual conflict. We found that regardless of the female type (standard vs. nutritionally impoverished), spatial structuring has a significant negative impact on fitness components in females (but not in males) – possibly indicating the cost of dispersal. However, contrary to the expectations from the hitherto reported complex environment effect on sexual conflict wherein spatial complexity is expected to reduce ISC and the resulting gender load (Yun et al., 2017; Malek & Long 2019), we found no evidence suggesting a reduction in ISC in structured habitat. However, we found the cost of male exposure to be absent for poor condition females. We discuss the broader implications of our findings in light of existing understanding of sexual conflict, dispersal and life history theories.

Yun et al., (2017) reported that physical complexity of the habitat can modulate the intensity of ISC - bringing down male-female interaction rate, giving females opportunity to seek refuges, and even changing the outcome of male mate choice for high fecundity females. This and the subsequent follow-up study from the same group was an important step towards bridging the intellectual gap between sexual conflict research and the nuanced ecology of a natural population. We investigated another component of a natural population that is usually simplified in laboratory microcosm setups - fragmented, discontinuous nature of a habitat. In our experimental setups, we found clear evidence of ISC, but unlike Yun et al., (2017), no evidence indicating a decline in ISC in structured habitat. Instead, we found significant detrimental effects of habitat type treatment on female progeny output as well as starvation resistance. Such negative effect of our habitat type treatment (viz., unstructured vs. structured) was unsurprising, and was a possibly a representation of hitherto reported cost of dispersal (Mishra et al., 2022). Though we did not record the degree of dispersal of individuals across the vials during our assay, we have ample data on the movement tendencies of flies under similar setups, and that using a two-patch system (see supplementary information for experimental details of dispersal tendency). A key observation in the two-patch pilot experiment was the higher tendency of females to spontaneously move between patches. Such female biased dispersal tendency is a perfectly compatible explanation for the observations in our Experiment 2, wherein only females, but not males, were found to show fitness costs of living in the fragmented habitat. Further, a three-patch system pilot-assay also suggested that females were more often spotted in the connecting corridors (i.e., the tubes connecting the three vials), further indicating that the females are on the move more often.

Theoretically, in a fragmented habitat, females can escape male congregation by moving to a patch with less male abundance. Interestingly, when Mishra et al., (2018b) found density dependent dispersal in female *D. melanogaster*, such density dependence was present only when the sexes were held together, and not when they were kept apart. These results seem to suggest a role of the presence of males on female dispersal, potentially invoking the mate harm avoidance hypothesis. Thus, in a habitat that allows such movement opportunities, females can be expected to escape mate harm, at least to some extent, by moving to a neighbouring patch, potentially resulting in a reduction in population level intensity of ISC. However, such male-avoidance related female dispersal can result in temporary shortage of females in a patch or temporary increase in the intensity of male-male competition resulting in increased ‘mate-finding dispersal’ in males (Mishra et al., 2020). This can in turn result in the equalisation of sex ratio across habitat patches, ameliorating the advantage of female biased dispersal, if any. Indeed, we have observed a substantial temporal and spatial variation in sex ratio in our pilot assay, results of which have been included in the supplementary information. However, none of these behavioural patterns seemed have any impact on the intensity of sexual conflict in our experimental setup. It is quite possible that reduction in ISC under a complex environment, as reported by Yun et al.,(2017) and Malek & Long, (2018), critically depends on the availability of spatial refuge to the females. Therefore, unless spatial structuring also includes access to refuges, it would not lead to reduced ISC. However, Byrne and Rice, (2008) did not find any evidence of access to refuge leading to reduction in ISC. Importantly, in natural populations habitat structure is usually accompanied by a number of other ecological variables, such as, heterogeneity in resource availability, abundance of natural enemies. It is possible that the presence of such heterogeneity in cost and reward of different habitat patches have direct or indirect implications in determining the strength of ISC. For example, the detrimental effect of larger males of *Aquarius remigis* seems to be stronger in habitats with high risk of predation (Sih & Krupa, 1992). In the Trinidadian guppies, *Poecilia reticulata*, males in habitat patches with high risk of predation tend to be more persuasive towards females, resulting in increased mate harassment (Magurran & Seghers, 1994). Hence, in such a species both predation and habitat structure seem to inflate the effect of ISC, which can in turn have important consequences on dispersal across habitat patches (for example, see Rowe et al., 1994).

Female-biased dispersal has been previously reported in several animals (Iliadi et al., 2002, Mishra et al., 2018a, b). As it turns out, female-biased dispersal is fairly common in birds (Clarke et al., 1997), and also seen in mammals (Favre et al., 1997), fishes (Yue et al., 2012) and invertebrates (Caudill, 2003; Sundström et al., 2003; Beirinckx et al., 2006). It is more commonly associated with competition for resources instead of mates, which is often a driver of male biased dispersal (Li & Kokko, 2019). An interesting exercise is to consider the consequences of female philopatry, the tendency to remain in the patch of birth. There are two main costs of philopatry for *D. melanogaster* females, which live on ephemeral food patches - (a) increasing competition for food (Bath et al., 2018), which is a primary determinant of female reproductive output, and (b) increasingly unfavourable environment for offspring development (Botella et al., 1985). Incidentally, these are the two main components of female fitness.

Male fitness, on the other hand, depends primarily on mating success (Bateman, 1948), and all else being equal, it is much harder for males to increase their chances of mating by simply dispersing to a different patch, unless the resident patch turns out to be mate deprived (Joshi et al., 1999). Thus, it is reasonable to suggest that a female biased dispersal rate is perhaps an adaptive demographic outcome given the behaviour and ecology of *D. melanogaster*. However, as our results suggest, females suffer a significant negative impact of living in a fragmented habitat - possibly due to costs of dispersal. In the long run, this should strengthen natural selection acting on females. Though our results rule out condition-dependence of sex-biased dispersal tendency (see supplementary information), it will be interesting to find out if such sex biased dispersal persists under a variety of other conditions. There is already some evidence that population density can break down this pattern (Mishra et al., 2018b).

Counterintuitively, reduction in progeny production on extended male exposure (i.e., LE vs. CE comparison) was found to be nonsignificant for poor quality females (i.e., females raised on a nutritionally impoverished diet), whereas the same was significant for standard females. This could be due to (a) poor condition females being closer to the fitness lower bound (i.e., zero or very low progeny output) – making the detection of fitness reduction upon prolonged male exposure much harder to detect, or (b) poor condition females being actually harassed less by the males. The latter is possible if males had preference against poor condition females. Male mate choice in *D. melanogaster* is well documented, and indicates adaptive choice towards female qualities associated with higher fecundity (Byrne & Rice, 2006; Nandy et al., 2012; Edward & Chapman, 2013; Arbuthnott et al., 2017). Long et al., (2009) showed non-virgin females of higher quality (viz., larger females) to receive greater male courtship compared to that received by low quality females. Such differential courtship resulted in high quality females suffering higher cost of male exposure (Long et al., 2009). Though our results fit this theory, we currently lack data required for a more direct test. To test the first possibility, we measured relative reduction (RR) score, a matric designed to capture the fitness cost of male encounter in females, controlling for the basal female fitness (female fitness under LE condition). Details of RR score and the analysis can be found in supplementary information (Table S7). Briefly, RR score is the ratio of fitness difference between the LE and CE conditions, and the fitness under LE condition. Analysis of RR score suggested that the effect of female type on RR score was non-significant –indicating that the degree of fitness reduction was not different across the female types. Hence, the significant ‘female type × male exposure’ interaction that we found in the Experiment 2 was possibly because of the fitness floor effect mentioned above. Hence, it is better to take caution interpreting the results of our analysis.

Interestingly, the analysis further indicated a significant effect of habitat type on RR score, such that it was higher in structured habitat type. Hence, the RR score analysis indicates that habitat type does impact the ISC outcome. Measured by the RR score, structured habitat seems to have higher ISC. However, the power of this analysis is limited since it relies only on a total of 12 data points. Hence, we do not intend to draw any general conclusion based on this (see supplementary information).

In conclusion, our results are, to the best of our knowledge, the first empirical test of the effect of spatial structure on the population level of interlocus sexual conflict. Our results demonstrate that spatial structuring by itself may not be sufficient to reduce the ISC unlike complexity of environment. Habitat structure can, however, impose additional fitness costs if it results in inter-patch dispersal, and dispersive behaviour itself is costly. Such a fitness cost can be sex specific, if there is sex difference in dispersal tendency. Future investigations should assess the generality of these findings under variability of different components of environment, including patch quality, predation risk, and intensity of competition.

## Conclusion

Interlocus sexual conflict, wherein male coercion including courtship and mating attempts reduce female fitness, is a widely observed attribute many animals. Theories predict spatial structure of a population modulating the intensity of such conflict by allowing females escape male coercion, yet there is little empirical support. Our experimental study using fruit flies show that habitat structure does not necessarily reduce conflict, but may impose additional sex-specific fitness cost due to energetically expensive sex-biased dispersal between patches.

