## Supplementary information for "Spatial structure imposes sex-specific costs but does not reduce interlocus sexual conflict"

**Supplementary material**

**A. Experimental structured habitat and the dispersal dynamics therein**

For the purpose of the assays (mentioned in the main text) we have used a simple three-patch habitat setup as our experimental structured habitat. These setups were created by connecting three culture vials by narrow tubes (diameter 0.6 cm) of 12.5 cm length that allow flies to move between vials. These tubes allow movement both directions. These setups were used as our experimental fragmented interconnected habitat (Figure S1a), whereas unfragmented habitat was three standard culture vials without connection between them (Figure S1b).

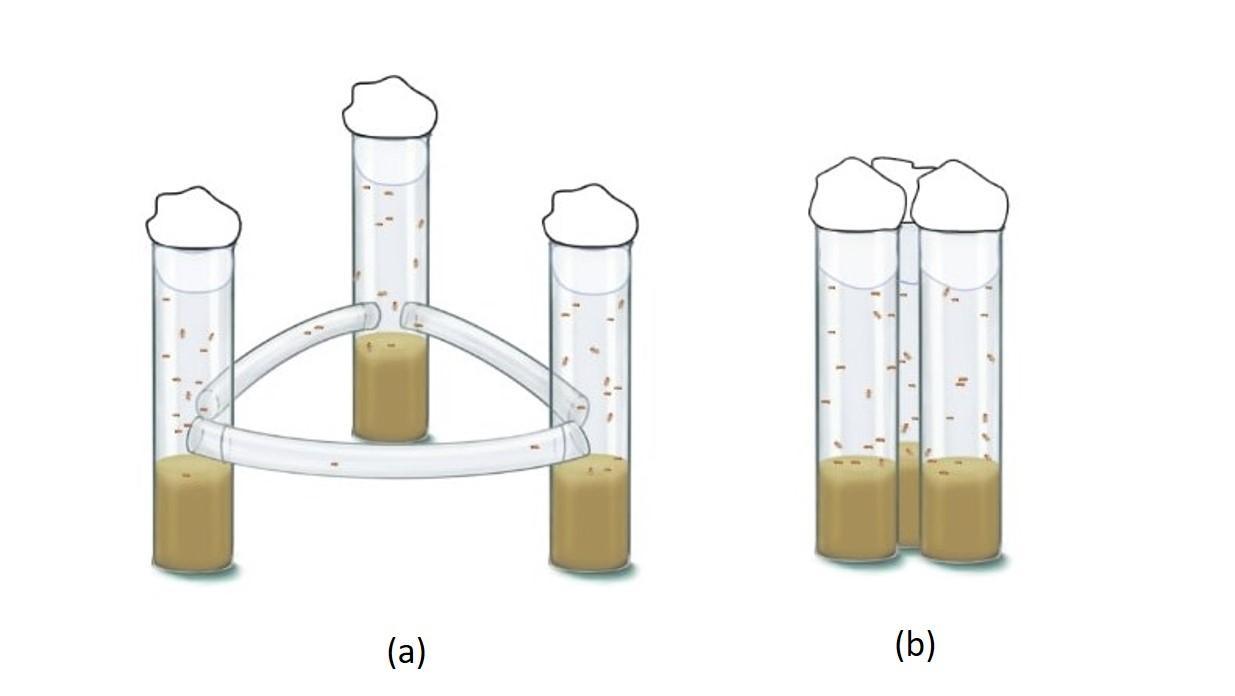

**Figure S1: Schematic representation of the two experimental habitat types.** In (a) Structured, and (b) Unstructured habitat setup types depicted in the figure, circles represent standard food vials (dimension: 25mm diameter × 90mm height). In Structured setup, the double headed arrows represent the interconnecting tubes (corridors) of length 12.5 cm, and diameter 0.6 cm.

To understand the dispersal dynamics, i.e., movement of individuals across the three patches in the experimental structured habitat setup, a pilot assay was conducted where observation with 10 replicates of such structured habitat setup recorded. The assay was set up with an initial condition wherein nine individuals (4 females and 5 males) were introduced in each patch. Thereafter, the numbers of males and females in different parts of the setup (i.e., patch1-3, and the connecting tubes) were recorded at regular intervals (see Fig S2 for the interval identity) for 48 hours.

Fig S2 shows the observed temporal variation across the three patches within a setup in total number of flies and proportion of females (Fig S2b). Figure S2a shows the variance (±95% CI) in number individuals per patch at different time intervals starting from the onset of the assay. Figure S2b shows the same for sex ratio (measured as proportion of females). As can be seen in these figures, there was a substantial spatial (across patches) variance in both these demographic parameters, and temporal (across time points) variation is such variation itself. These variations indicate potential change in intra and inter-sexual interactions in our experimental structured habitat.

Further, a substantial number of flies were also observed in the connecting tubes (i.e., the corridor connecting the patches) at almost all observation points. Interestingly, females were found significantly more often than males in such cases (Mean number $\pm$95% CI: Female: 21.9 $\pm$4.96, Male: 7 $\pm$1.54 Kruskal-Wallis rank sum test, $\chi^{2}$= 13.86, df = 1, p < 0.01, Figure S3).

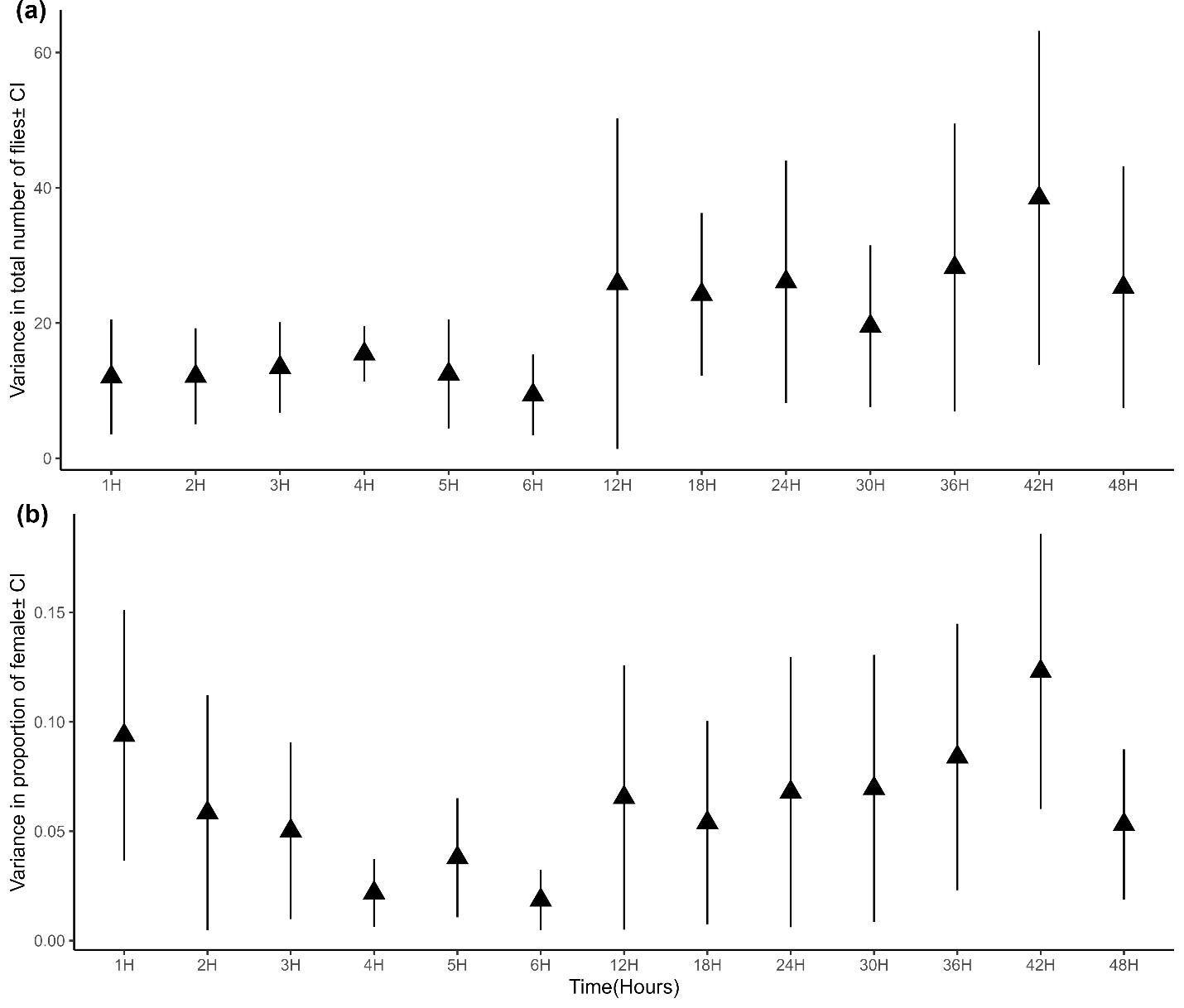

**Figure S2: Variation in number of flies and sex ratio across patches.** Starting from 1 hour following the onset of the assay, number of males and females in each vial (i.e., each patch in the three-patch setup) was counted. The starting conditions were nine (4 females and 5 males) individuals in each vial – therefore, a sex ratio of female: male = 4:5 (i.e., proportion of females is 0.44). (a) Variance in number of flies per vial was calculated for each replicate setup. A 95% confidence interval (CI) was computed using the variance value from all ten replicate setups. The plot shows such variance ± 95% CI in number of flies at 13 observation points within the observation window of 48 hours. (b) Similarly, variance ± 95% CI was computed and plotted for sex ratio, where sex ratio is measured as proportion of females.

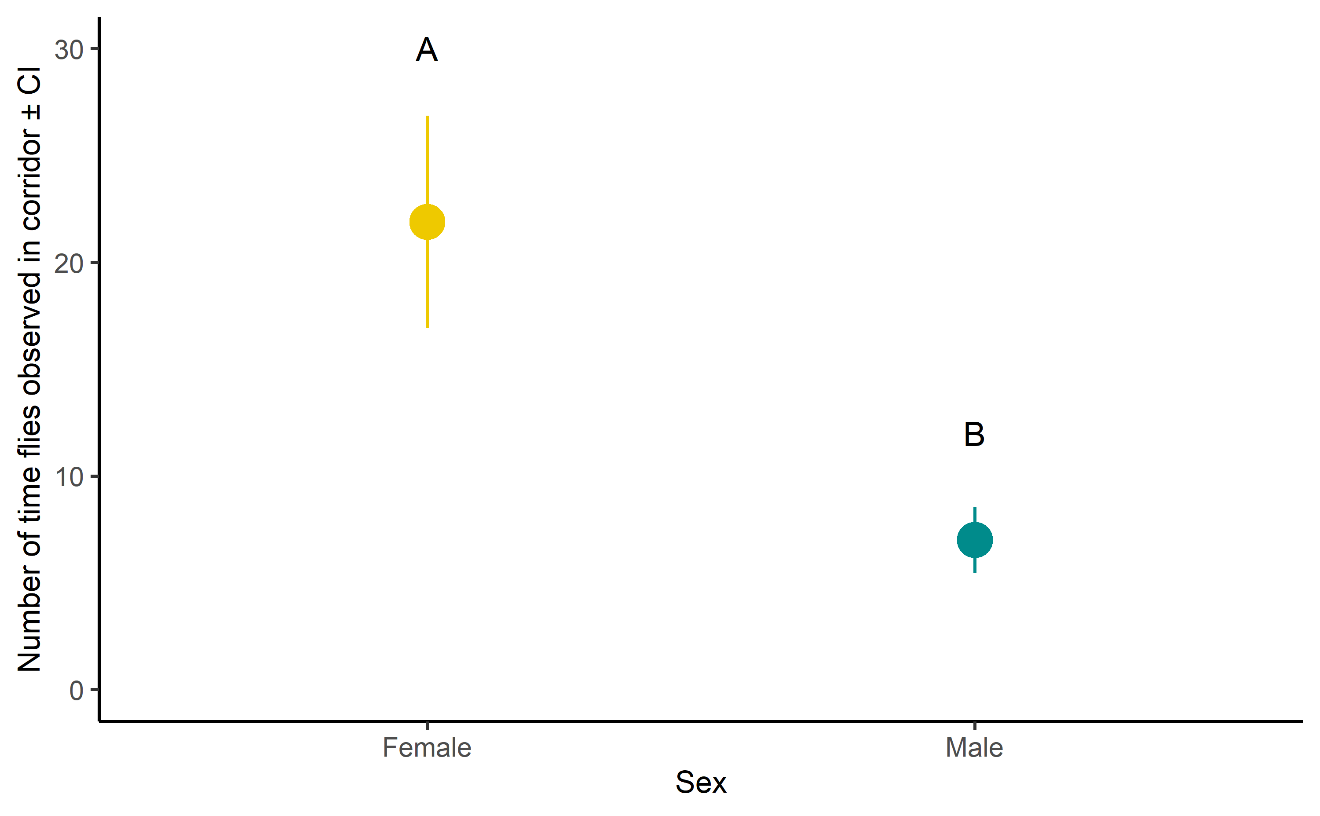

**Figure S3: Number of females and males in the corridor** (i.e., the tube connecting the vials in the setup) was counted at different time points. The plot shows the mean number of females and males across all observations and replicate setups. As suggested by the analysis, females were spotted significantly more often in corridors than males (Mean number $\pm$95% CI: Female: 21.9 $\pm$ 4.96, Male: 7 $\pm1.54,$ Kruskal-Wallis rank sum test, p-value<0.01). Treatments not sharing common alphabets are significantly different.

**B. Sex specificity in spontaneous dispersal: A two-patch system observation**

Eggs were collected from a randomly chosen population (the assay was done with BL_3,_ BL_4_, BL_5_), and cultured in standard vials at a density of 70 per vial with ~8ml food. The adults were allowed to emerge in these vials, upon which they were transferred to fresh food vials with standard food. On day12 from the egg-collection, i.e., at 2-3 days post-eclosion, two-patch dispersal assays were set up to measure dispersal tendencies of the experimental flies.

The dispersal assay setups used here were two-patch systems constructed by connecting a culture vial (with food) to a standard Drosophila culture bottle (Laxbro) with a tygon tube (Tarsons) of 0.6 cm inner diameter and 31cm length (see Fig. S4). The vial was used as the source patch, where flies were introduced to start the assay, while the bottle is used as the sink. To ensure unidirectional movement of the flies, the bottle hole is fitted with a pipette tip, cut in such a way that it allowed flies to exit the tube through it and enter the bottle. Observations in our laboratory showed that without the pipette tip arrangement, flies moved in both directions unabated (i.e., vial to bottle, as well as bottle to vial). However, this setup allowed only source to sink movement, but stopped all movement in the reverse direction.

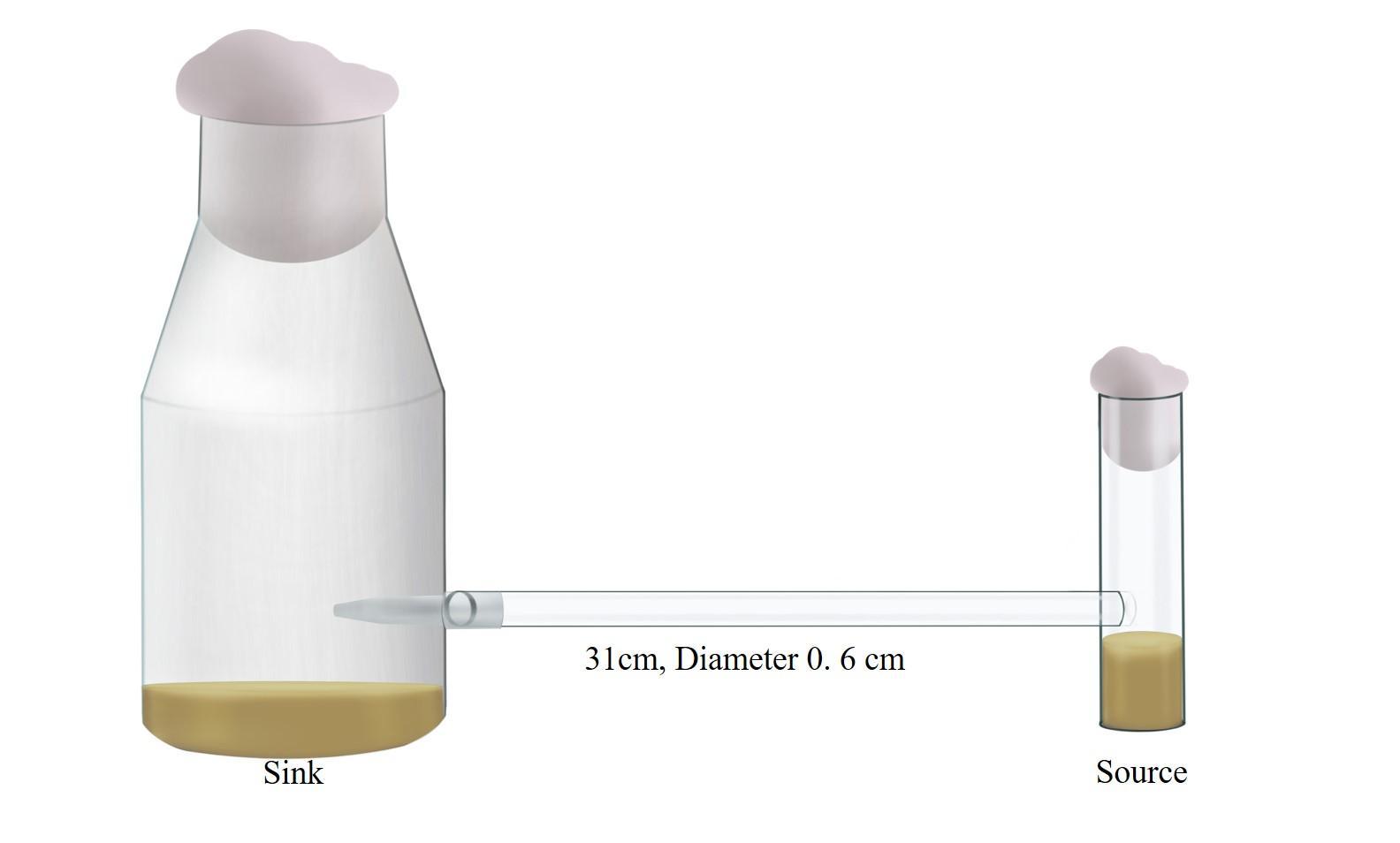

**Figure S4: A two-patch setup for dispersal assay**. A culture food vial (Source patch) connected to a standard Drosophila culture bottle (Sink patch) by a tube of 0.6 cm inner diameter and 31cm length.

To start the assay, the 15 males and 15 females were introduced in a source patch (i.e., the vial) by using light CO_2_-anaesthesia. The setup was then placed horizontally in a well-lit place with illumination on all four sides. These setups were randomly placed to avoid any positional bias. They were left undisturbed for six hours for observation. Number of flies reaching the sink patch (i.e., the bottles) was counted after every two hours, until six hours from the start of the assay. Flies in the sink patch were sexed and counted. Dispersal propensity (tendency to leave source patch) was measured as the proportion-moved for each sex at the end of 6 hours. Based on our preliminary observations, by 6 hours, on an average more than 50% females move away from the source patch whereas males hardly move. To avoid the confounding effect of changing sex ratio, we considered cumulative dispersal unit 6 hours for our analysis. The entire experiment was conducted in three random blocks using BL_3_, BL_4_ and BL_5_ populations with 12 replicates per Block.

Dispersal propensity was analysed using General Linear Mixed Model (GLMM) - modelled with binomial family distribution and logit link function treating Sex as fixed effect, while block and interactions involving block as random factors using package lme4 and function glmer with binomial family distribution and logit link in R (R Version 4.2.0). Analysis of dispersal propensity showed significant effect of Sex, and Sex × Block interaction. Therefore, we re-analysed dispersal propensity for each block. This was done using GLM, with Sex as the only fixed factor. Sex was found to have significant effect on dispersal tendency in all three blocks (Table S1). Interestingly, females have significantly higher dispersal tendency than males in all the blocks (Figure S5). Hence, our results implied a significant female biased dispersal tendency. Incidentally, this observation also explains spotting of females more than males in the connecting corridors of the three-patch setups (see Figure S3).

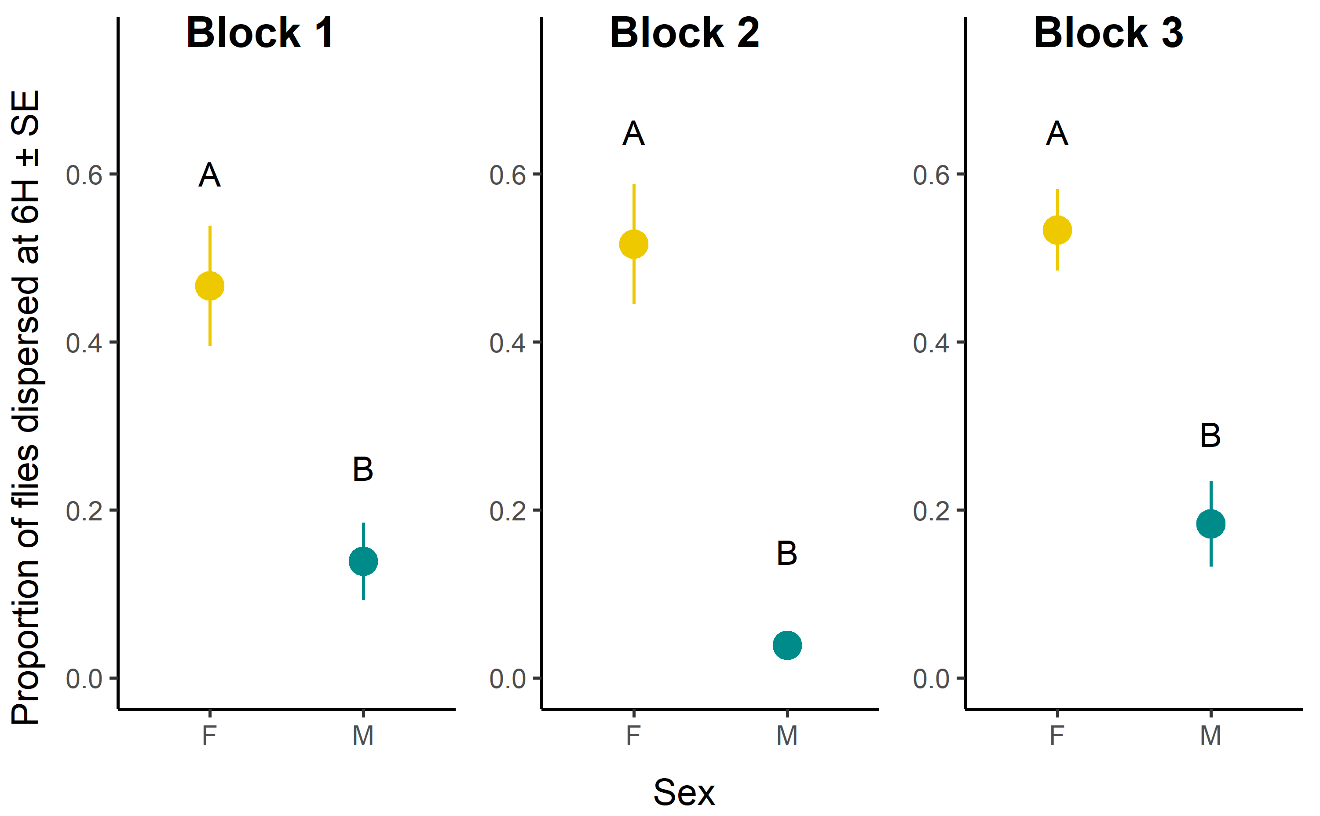

**Figure S5: Results from Experiment 1. Effect of sex on dispersal pattern**. Proportion of females (denoted with yellow color) and males (denoted with cyan color), within a replicate two-patch setup, that dispersed from the source to sink by 6hour duration is considered as a measure of dispersal propensity of that replicate and was used as the unit of analysis. The plots (a), (b) and (c) represents data from Block-1, Block-2 and Block-3 respectively. Treatments not sharing common alphabets are significantly different.

**Table S1: Summary of the analysis of Experiment 1, results from dispersal propensity assay.** The dispersal tendency results were analysed for each Block separately using a general linear mixed-effect model (GLMM) where Sex was fitted as fixed factor. Dispersal propensity was calculated as proportion-dispersed from source to sink for each sex at 6H from introduction of flies to source patch. Values from each dispersal replicate setup were used as the unit of analysis. Statistically significant p-values are mentioned in bold font style.

| **Contrast** | Blocks | ${}Chi sq$ | **Df** | **p-value** |
| --- | --- | --- | --- | --- |
| Sex | Block 1 | 26.27 | 1 | **<0.01** |
|  | Block 2 | 71.72 | 1 | **<0.01** |
|  | Block 3 | 23.98 | 1 | **<0.01** |

**Table S2: Food composition and recipe.** Control (i.e., complete diet), and 20%, 40%, 60% dilution of the nutritional components of the standard food.

| **Ingredients** | **Control** | **20% diluted** | **40% diluted** | **60% diluted** |
| --- | --- | --- | --- | --- |
| Banana (gm) | 205 | 164 | 123 | 82 |
| Barley (gm) | 25 | 20 | 15 | 10 |
| Jaggery (gm) | 35 | 28 | 21 | 14 |
| Yeast (gm) | 36 | 28.8 | 21.6 | 14.4 |
| Agar-agar (gm) | 12.4 | 12.4 | 12.4 | 12.4 |
| Water (ml)  [to mix with agar-agar] | 1000 | 1000 | 1000 | 1000 |
| Ethanol (ml)  [to mix with yeast] | 22 | 22 | 22 | 22 |
| Water (ml) | 180 | 180 | 180 | 180 |
| p-Hydroxy methyl benzoate | 2.4 | 2.4 | 2.4 | 2.4 |
| Ethanol (ml)  [to mix with benzoate] | 23 | 23 | 23 | 23 |

**C. Effect of dietary treatment on dispersal tendency**

To this effect, we raised flies on a range of dietary conditions - control (i.e., complete diet), and 20%, 40%, 60% dilution of the nutritional components of the standard food (see Table S2, for the detail regarding food composition and recipe).

To start the assay, the 15 males and 15 females were introduced in a source patch (i.e., the vial) by using light CO_2_-anaesthesia. The setup was then placed horizontally in a well-lit place with illumination on all four sides. For each diet regime, we set up 12 replicate sets. These setups were randomly placed to avoid any positional bias. They were left undisturbed and every two hours and for six hours the number of the flies in the sink patch (i.e., the bottles) were sexed and counted. Dispersal tendency was measured by calculating the proportion of females moved in sink at the end of 6 hours This experiment was also conducted in three random blocks using BL_3_, BL_4_ and BL_5_ populations.

Dispersal tendency was analysed using General Linear Mixed Model and modelled with binomial family distribution and logit link, using dietary condition (treatment) and sex as fixed effects, and block as random factor using package lme4, and function glmer in R version 4.2.0

Analysis of dispersal tendency showed a significant effect of sex. As usual, females disperse significantly more than males. However, we did not find a significant effect of treatment (see Table S3, Figure S6). Based on these results, we used 40% dilution treatment to generate the poor-condition females in our Experiment 2 to avoid variation in dispersal tendency as a confounding factor.

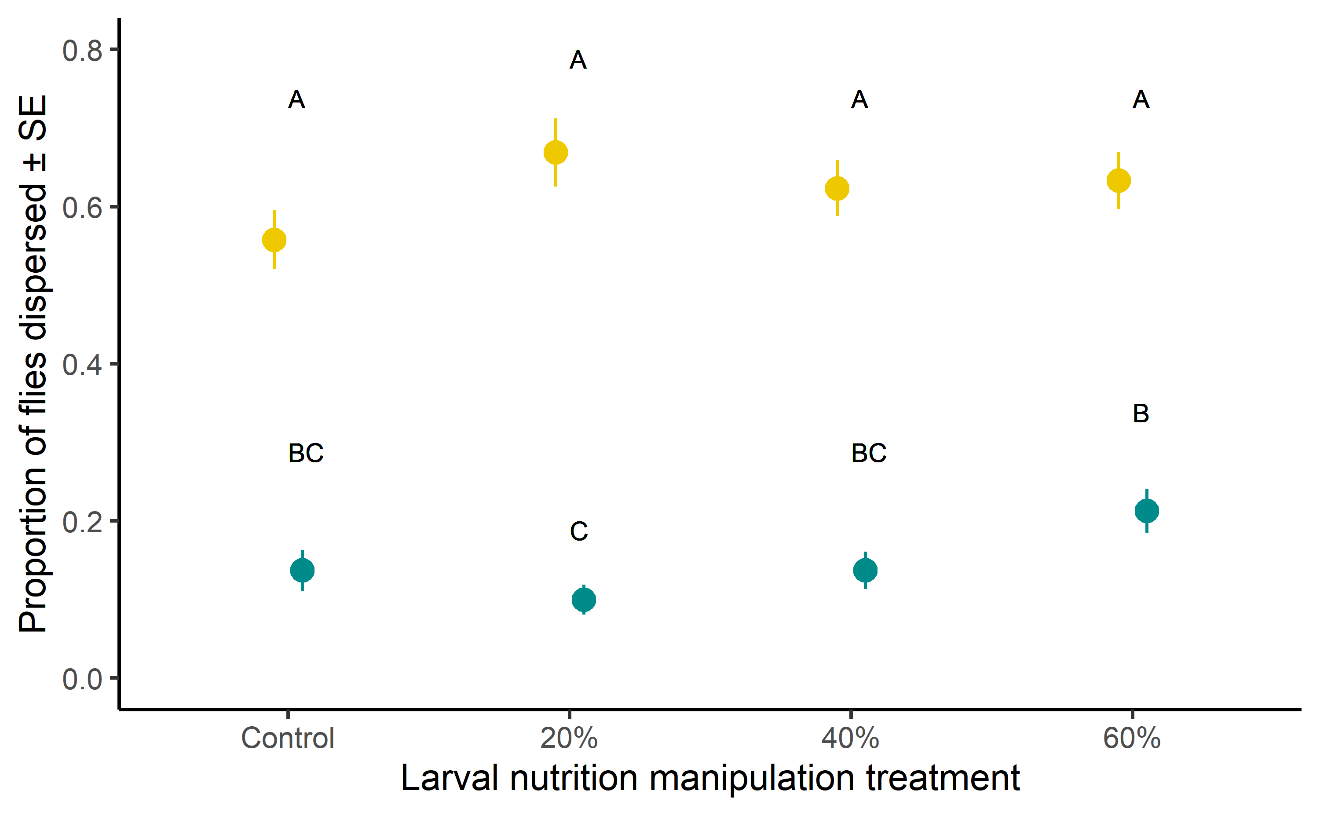

**Figure S6: Results from Experiment 2. Effect of diet treatment on dispersal tendency**. Larval nutrition manipulation treatment represents alteration of the nutritional quality of the food wherein control represents the standard banana-jaggery-barley-yeast food, and 20%, 40%, 60% were the diluted food where only the nutritional components were diluted by the corresponding amount. Proportion of females (denoted with yellow colour) and males (denoted with cyan colour), within a replicate two-patch setup, that dispersed from the source to sink following a six-hour duration is considered as a measure of dispersal tendency of that replicate, and was used as the unit of analysis. The plot represents data combining all three blocks. Treatments not sharing common alphabets are significantly different.

**Table S3: Summary of the analysis of results from dispersal tendency assay.** The dispersal tendency results were analysed using a general linear mixed-effect model (GLMM) where treatment (diet condition: control, 20%, 40%, and 60% dilutions of control) and Sex were fitted as fixed factors. Block as random factors. Dispersal tendency was calculated as proportion-dispersed from source to sink for each sex. Values from each dispersal replicate setup were used as the unit of analysis. Statistically significant p-values are mentioned in bold font style.

| **Contrast** | $\chi^{2}$ | **DF** | **p-value** |
| --- | --- | --- | --- |
| Treatment | 6.31 | 3 | 0.09 |
| Sex | 320.96 | 1 | **<0.01** |
| Treatment× Sex | 15.33 | 3 | **<0.01** |

**Table S4: Results of pairwise comparisons on progeny count data from Experiment 1.** showing a significant difference of progeny count in CE treatment and LE treatment only for standard females (mean ± se, Standard CE: 6.18 ±1.39, Standard LE: 10.86 ±1.39), but not for poor females raised in poor diet (mean ± se, Poor CE: 3.91 ±1.39, Poor LE: 5.81 ±1.40).

| **Contrast** | **estimate** | **SE** | **df** | **t.ratio** | **p.value** |
| --- | --- | --- | --- | --- | --- |
| Poor CE - Standard CE | -2.27 | 0.93 | 3.19 | -2.42 | 0.24 |
| Poor CE - Poor LE | -1.90 | 0.86 | 3.52 | -2.20 | 0.27 |
| Poor CE - Standard LE | -6.94 | 1.08 | 3.92 | -6.42 | **0.01** |
| Standard CE - Poor LE | 0.36 | 1.08 | 3.96 | 0.34 | 0.98 |
| Standard CE - Standard LE | -4.67 | 0.85 | 3.38 | -5.49 | **0.02** |
| Poor LE - Standard LE | 5.04 | 0.94 | 3.22 | -5.36 | **0.03** |

**Table S5: Model structures of the different analyses in the Experiment 1 and 2.** The actual codes for each analysis are mention. The abbreviations used are as follows: MPO = Mean progeny output, M.Exposure: Male exposure type, MST = Mean survival time. “Habitat” and “Female” refer to the habitat type and female type treatments in the corresponding experiment.

| **Experiment 1** | |
| --- | --- |
| Progeny count | MPO ~ M.Exposure * Habitat + (1\|BLOCK) + (1\|BLOCK:M.Exposure) + (1\|BLOCK:Habitat) + (1\|BLOCK:Habitat:M.Exposure) |
| Starvation survival time | MST ~ M.Exposure * Habitat + (1\|BLOCK) + (1\|BLOCK:M.Exposure) + (1\|BLOCK:Habitat) + (1\|BLOCK:Habitat:M.Exposure) |
| **Experiment 2** | |
| Progeny count | MPO ~ Female * M.Exposure * Habitat + (1\|BLOCK) + (1\|BLOCK:Female) + (1\|BLOCK:M.Exposure) + (1\|BLOCK:Habitat) + (1\|BLOCK:Female:M.Exposure) + (1\|BLOCK:Habitat:M.Exposure) + (1\|BLOCK:Habitat:Female) + (1\|BLOCK:Female:Habitat:M.Exposure) |
| Starvation survival time (Female) | MST~Female*M.Exposure*Habitat+(1\|BLOCK)+(1\|BLOCK:Female)+(1\|BLOCK:M.Exposure)+(1\|BLOCK:Habitat)+(1\|BLOCK:Female:M.Exposure)+(1\|BLOCK:Habitat:M.Exposure)+(1\|BLOCK:Habitat:Female)+(1\|BLOCK:Female:Habitat:M.Exposure) |
| Starvation survival time (Male) | MST~Female*Habitat+(1\|BLOCK)+(1\|BLOCK:Female)+(1\|BLOCK:Habitat)+(1\|BLOCK:Habitat:Female ) |

**Table S6:** Effect of random blocks and their interactions with the fixed factors for Progeny count and Starvation survival time

| **Trait** | **Effects** | **npar** | **AIC** | **logLik** | **deviance** | **Chisq** | **DF** | **Pr(>Chisq)** |
| --- | --- | --- | --- | --- | --- | --- | --- | --- |
| **Experiment 1** | | | | | | | | |
| Progeny  count | Block$\times$Male exposure | 8 | 547.94 | -265.97 | 531.94 | 0 | 1 | 1 |
|  | Block$\times$Habitat type | 8 | 547.94 | -265.97 | 531.94 | 0 | 1 | 1 |
|  | Block$\times$Male exposure $\times$ Habitat type | 8 | 547.94 | -265.97 | 531.94 | 0 | 1 | 1 |
| Starvation survival time | Block$\times$Male exposure | 8 | 989.13 | -486.57 | 973.13 | 0 | 1 | 1 |
|  | Block$\times$Habitat type | 8 | 989.13 | -486.57 | 973.13 | 0 | 1 | 1 |
|  | Block$\times$Male exposure $\times$ Habitat type | 8 | 989.13 | -486.57 | 973.13 | 0 | 1 | 1 |
| **Experiment 2** | | | | | | | | |
| Progeny  count | Block$\times$Female type | 16 | 1018.0 | -493.03 | 986.05 | 0.00 | 1 | 1 |
|  | Block$\times$Male exposure | 16 | 1018.0 | -493.03 | 986.05 | 0.00 | 1 | 1 |
|  | Block$\times$Habitat type | 16 | 1018.0 | -493.03 | 986.05 | 0.00 | 1 | 1 |
|  | Block$\times$Female $\times$Male exposure | 16 | 1018.0 | -493.03 | 986.05 | 0.00 | 1 | 1 |
|  | Block$\times$Female Type$\times$ Habitat type | 16 | 1018.0 | -493.03 | 986.05 | 0.00 | 1 | 1 |
|  | Block$\times$Male exposure $\times$ Habitat type | 16 | 1018.0 | -493.03 | 986.05 | 0.00 | 1 | 1 |
|  | Block$\times$Female type $\times$ Male exposure$\times$ Habitat type | 16 | 1018.0 | -493.03 | 986.05 | 0.00 | 1 | 1 |
| Starvation survival time  (Female) | Block$\times$Female type | 16 | 1418 | -698 | 1396 | 0.00 | 1 | 1 |
|  | Block$\times$Male exposure | 16 | 1418 | -698 | 1396 | 0.00 | 1 | 1 |
|  | Block$\times$Habitat type | 16 | 1418 | -698 | 1396 | 0.00 | 1 | 1 |
|  | Block$\times$Female type $\times$ Male exposure | 16 | 1418 | -698 | 1396 | 0.00 | 1 | 1 |
|  | Block$\times$Female type $\times$ Habitat type | 16 | 1418 | -698 | 1396 | 0.00 | 1 | 1 |
|  | Block$\times$Male exposure $\times$ Habitat type | 16 | 1418 | -698 | 1396 | 0.00 | 1 | 1 |
|  | Block$\times$Female type $\times$ Male exposure$\times$ Habitat type | 16 | 1418 | -698 | 1396 | 0.00 | 1 | 1 |
| Starvation  survival  time  (Male) | Block$\times$Female type | 8 | 833.26 | -408.63 | 817.26 | 1.96 | 1 | 0.16 |
|  | Block$\times$Habitat type | 8 | 831.91 | -407.96 | 815.91 | 0.61 | 1 | 0.43 |
|  | Block$\times$Female type $\times$ Habitat type | 8 | 831.3 | -407.65 | 815.3 | 0 | 1 | 1 |

**D. Assessment of the cost of male encounter through Relative Reduction in progeny production**

In our Experiment 2, we found a significant effect of female type × male exposure interaction on female progeny output (see Table 1). Standard females were more adversely affected by continued male exposure than the poor females in our assay. The poor condition females being closer to the fitness lower bound (i.e., zero or very low progeny output), it is possible that the fitness reduction upon prolonged male exposure is difficult to resolve by analysing simple differences in mean (e.g., between low and high male exposure). Therefore, we quantified relative reduction (called RR hereafter) in progeny output which accounts for the baseline fitness difference among females. We computed RR score for each assay population using the block

means in the following formula, and thus we had three RR values coming from three blocks:

$$RR= \frac{mean progeny count in LE-mean progeny count in CE}{mean progeny count in LE}$$

The RR score was then analysed using mixed-model analysis of variance without replicate in Statistica (version 13.3, TIBCO). Block was treated as a random factor, while female type and habitat type were treated as fixed factors. Nonsignificant effect of female on RR score suggests that relative reduction in progeny output did not differ between two types of female and significant female type × male exposure interaction in experiment 2 was indeed because of the floor effect and therefore an artifact of statistical analysis. Nonsignificant female × habitat confirms that the relative reduction in fitness across two females was also unaffected by habitat type (Figure S7).

**Table S7:** Effect of female type and habitat type on relativised reduction in progeny output (RR score). Significant p-value is shown in bold.

| **Contrast** | **DF** | **MS** | **Den Syn Error df** | **Den Syn Error MS** | **F** | **p** |
| --- | --- | --- | --- | --- | --- | --- |
| Female | 1 | 0.02 | 2.00 | .00 | 3.29 | 0.21 |
| Habitat | 1 | 0.10 | 2.00 | .00 | 30.14 | **0.03** |
| Female × Habitat | 1 | 0.06 | 2.00 | .03 | 1.99 | 0.29 |

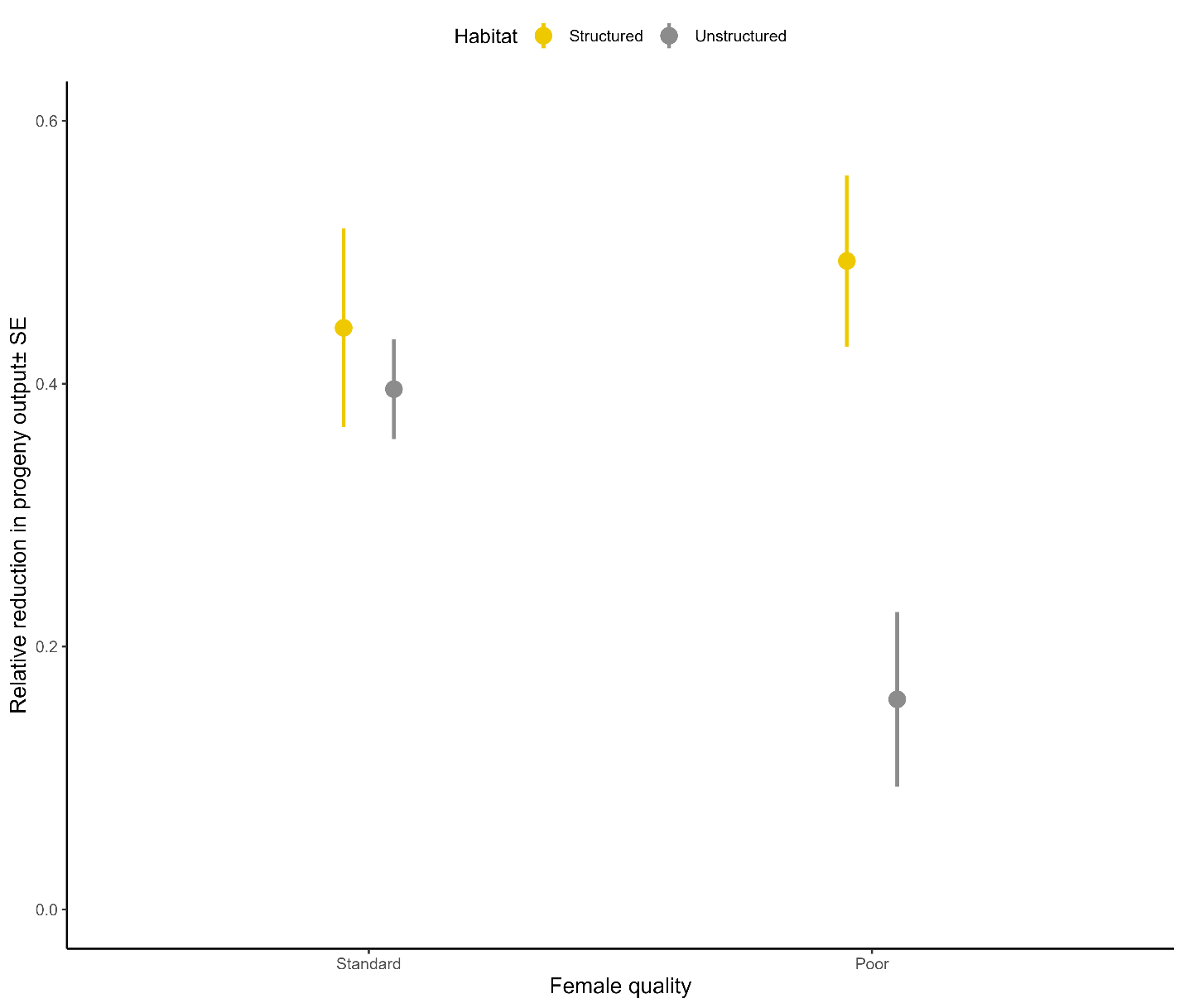

**Figure S7: The Relative reduction (RR) scores from Experiment 2.** RR scores were calculated for each block, using the block means of progeny count results. It measures the reduction in progeny production (and hence, fitness) on prolonged exposure to males (limited vs. continued exposure), while considering progeny output under limited male exposure to be an intrinsic progeny production ability. Hence, RR is a measure of cost of male exposure to the females. Effect if habitat type treatment for poor and standard females are shown and error bars represent standard error.
